# Tracking the cells of tumor origin in breast organoids by light sheet microscopy

**DOI:** 10.1101/617837

**Authors:** A Alladin, L Chaible, S Reither, M Löschinger, M Wachsmuth, JK Hériché, C Tischer, M Jechlinger

**Author notes:** Contributed equally (A Alladin, L Chaible). to whom correspondence should be sent.

## Abstract

How tumors arise from individual transformed cells within an intact epithelium is a central, yet unanswered question. Here, we developed a new methodology that combines breast tissue organoids, where oncogenes can be switched on in single cells, with light-sheet imaging that allows us to track cell fates using a big-image-data analysis workflow. The power of this integrated approach is illustrated by our finding that small local groups of transformed cells form tumors while isolated transformed cells do not.

## Main Text

Organoid cultures grown from cell-lines or primary cells have been successfully employed to study molecular mechanisms during different stages of tumorigenesis^1–3^. However, they usually allow oncogenic activation in all cells of the tissue and therefore cannot reproduce the localized transformation at a defined part of the tissue that is seen in the patient situation. Hence, stochastic tumor models need to be established wherein only a few tumorigenic cells expand in a normal epithelium. It is of key importance to visualize the interactions between a transformed cell and the normal neighboring epithelium in order to better understand the tumor initiation process in the context of its immediate microenvironment^4^.

Long term imaging of complex primary organoids has been achieved via light-sheet microscopy^5, 6^ and have benefited from the lower phototoxicity. However, past advancements still came with trade-offs resulting in limited cellular and temporal resolution. Tracking single cell dynamics necessitates high resolution imaging which in turn limits the time frame in which organoids can be imaged without phototoxic effects^7^. Conversely, imaging primary organoids for longer time periods requires an offset of temporal and cellular resolution that eventually cannot allow single cell fate tracking^8^.

Here we present a novel stochastic model of breast tumorigenesis where only single cells express oncogenes in primary murine organoids. We thereby overcome the above-mentioned limitations of studying tumorgenesis events. Furthermore, we report long-term imaging of these organoids for the first time at a temporal resolution that allows us to follow single cell fates. We also integrate this approach with an image analysis pipeline capable of segmenting cells in their dynamic progression towards tumorigenesis, so they can be tracked individually over time.

For modelling tumorigenesis in breast tissue, we use an inducible model of breast cancer^9–11^ that has been shown to recapitulate hallmarks of human breast disease^3, 12^ (Fig.1a). In this tractable transgenic mouse model, the activity of two potent oncogenes -Myc and Neu (the rodent homolog for the human HER2 gene)- can be spatially limited by tissue specific expression of the rtTA inducer-protein to the cells of the mammary lineage and temporally controlled by the addition of doxycycline in the media or animal diet^13^. We adapt this tissue wide tumorigenesis model (tri-transgenic (T) model) to generate a stochastic system by retaining only the oncogenic constructs (bi-transgenic (B) model). The rtTA inducer gene is then lentivirally delivered to single cells, preventing tissue wide transformation.

**Figure 1.**
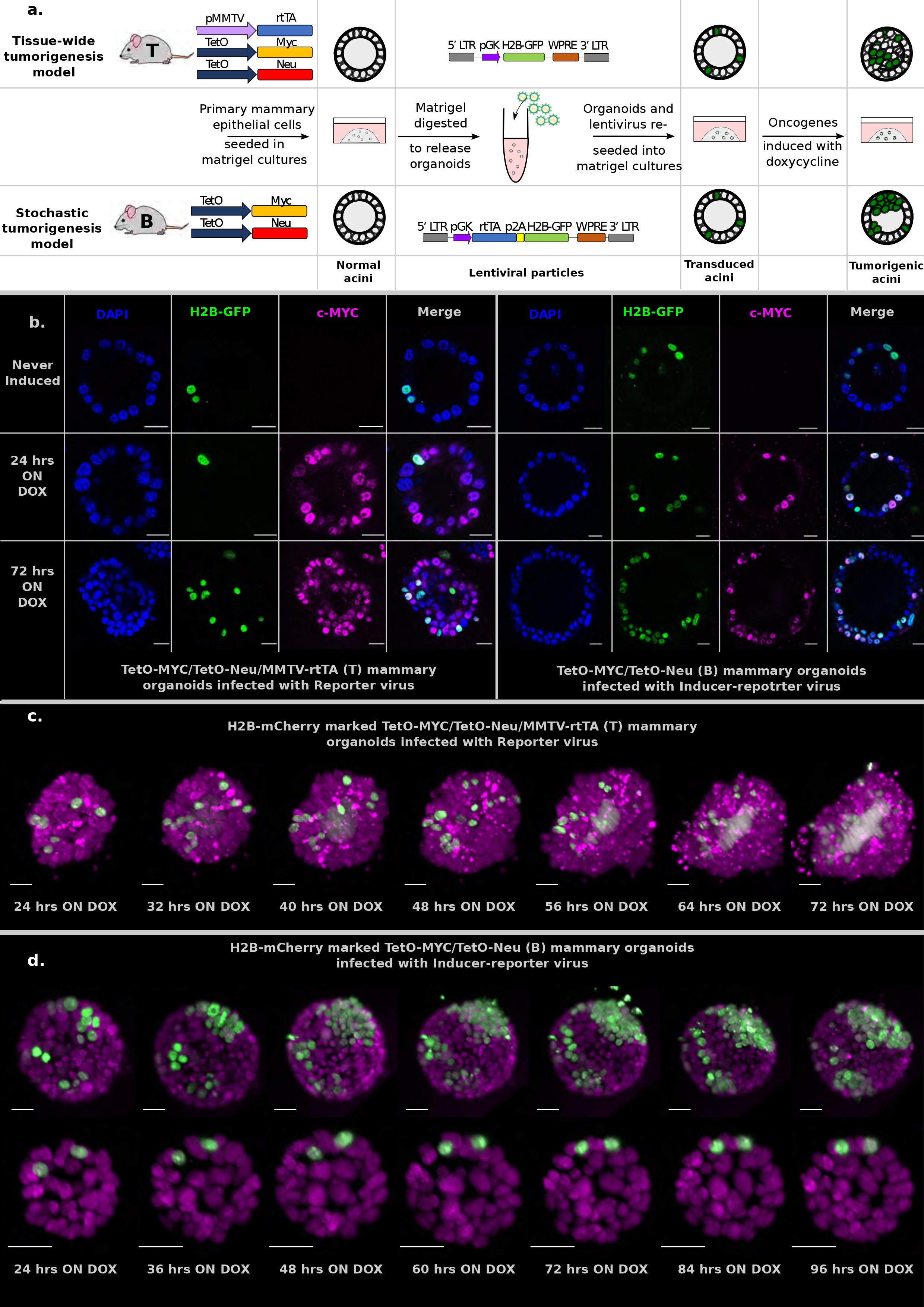
Characterization and imaging of stochastic tumorigenesis in mammary organoids. **(a)** Schematic representation of the mouse models and the *in vitro* culture methods used. Organoids are grown from single cells harvested from the mammary glands of either bi-trangenic (B) or tri-trangenic (T) mice, transduced with lentiviral particles in solution and re-seeded into 3D cultures. Doxycycline is added to the media to induce the expression of oncogenes in cells expressing rtTA. B mice have the c-MYC and Neu oncogene constructs in their genome. These oncogenes are activated in single cells infected with the Inducer-reporter (pLenti-rtTA-GFP) lentiviral particles, in the presence of doxycycline — modelling stochastic breast tumorigenesis (bottom panel). T mice have the rtTA transducer construct along with the oncogenes and all cells in T organoids can be induced to express oncogenes in 3D culture in the presence of doxycycline. T mice infected with Reporter (pLenti-NULL-GFP) lentiviral particles are used as infection controls (top panel). Both viral particles mark single cells in the organoids with H2B-GFP. **(b)** Representative immunoflourescence staining images of fixed 3D gels with B organoids transduced with Inducer-reporter virus or T organoids transduced with Reporter virus before induction (top), 24 hours post induction and (middle) and 72 hours post induction (bottom) with doxycycline. GFP expressing transduced cells (green), c-MYC oncogene (magenta), DAPI nuclear stain(blue). Scale bar, 10μm. **(c)** 3D images of selected timepoints during live-cell time-lapse microscopy of induced T organoids transduced with Reporter virus. GFP expressing transduced cells (green), c-MYC oncogene (magenta) **(d)** B organoids transduced with Inducer-reporter virus. GFP expressing transduced cells (green), c-MYC oncogene (magenta). Imaging was started 24 hours after oncogenic induction with doxycycline. The upper panel shows the proliferative phenotype seen with stochastic transformation, whereas the lower panel shows the non-proliferative phenotype observed in some stochastically transformed organoids. (Imaging conditions: H2B-mCherry 594nm Ex, 610 LP Em; H2B-GFP 488nm Ex and 497-554 nm Em). Scale bar, 20μm.

Primary mammary epithelial cells derived from transgenic mice were seeded in 3D matrigel as single cells, to form small acini. A small number of single cells in these acini were then transduced with lentiviral particles (Fig.1a, middle panel). In the tissue-wide tumorigenesis model(T), organoids were transduced with the reporter virus (pLv-pGK-H2B-GFP) that marks a subset of cells with H2B-GFP, while tissue wide rtTA expression is driven in all cells (Fig.1a, upper panel). To achieve stochastic tumorigenesis, bi-transgenic (B) organoids were transduced with the inducer-reporter virus (pLv-pGK-rtTA-p2A-H2B-GFP) that expresses rtTA and reporter H2B-GFP in only these single cells within the normal epithelium (Fig.1a, lower panel). Then, doxycycline was supplemented in the media to induce tumorigenic growth in rtTA expressing cells. Immunofluorescent staining of 3D matrigel cultures, for both sets of doxycycline-induced transduced organoids, was used to validate transgene specific protein expression of the c-MYC oncogene in only the transduced cells of B organoids as opposed to all cells of T organoids (Fig.1b). qPCR analysis of *Myc* and *Neu* mRNA expression was performed to normalize doxycycline dosage in both systems (Supplementary Fig.1).

Next, we bred the nuclear reporter H2B-mCherry into the T and B mice to mark all the cells in the organoids for inverted light-sheet microscopy (Luxendo InVi SPIM, Supplementary Fig.2). The InVI SPIM was adjusted for non-phototoxic, long-term imaging (up to 4 days, every 10 minutes with 1 μm z-spacing). The T/H2B-mCherry organoids transduced with reporter virus proliferated swiftly upon doxycycline addition showing expansion of both the marked and unmarked cells (Fig.1c); a sturdy tumor phenotype developed, manifested by multi-cell-layered rims and pronounced proliferation-associated-apoptosis in all organoids (Supplementary movie 1). In contrast, B/H2B-mCherry organoids transduced with inducer-reporter virus, displayed phenotypic variation upon induction of oncogenes in the transduced cells. Some organoids showed fast clonal expansion of oncogene-expressing cells that form multilayer clusters in the organoid rim. This proliferative phenotype seems to stem from several transduced cells in vicinity to each other at the start of time lapse imaging (Fig.1d, upper panel; Supplementary movie 2). Other, more sparsely infected organoids, did not sustain proliferation of the oncogene-expressing cells (Fig.1d, lower panel; Supplementary movie 3). Immunofluorescent staining 3D matrigel cultures for both sets of doxycycline-induced transduced organoids was performed to exclude imaging artefacts; consistent with the light-sheet movies, 3D gels grown in the incubator showed a similar dual phenotype for B organoids while T organoids consistently formed tumors upon oncogene induction (Supplementary Fig.3).

To analyze the dual-color light-sheet movies (H2B-mCherry—all cells in the organoid, H2B-GFP—transduced cells within the organoid) on a single cell level, we developed a big data compatible image analysis pipeline, using Fiji^14^ (plugins Big Data Processor^15^ and CATS^16^) and Imaris^17^, for efficient visualization of longitudinal image data, cell segmentation and tracking in 3D (Fig.2a, Supplementary Fig.4 and Fig.5). Cell tracking allowed us to follow the clonal evolution for each transduced cell in the organoid over 3 days. Single transduced cells within one organoid show a difference in proliferation and cell fate as indicated in representative tracks (Fig.2b).

**Figure 2.**
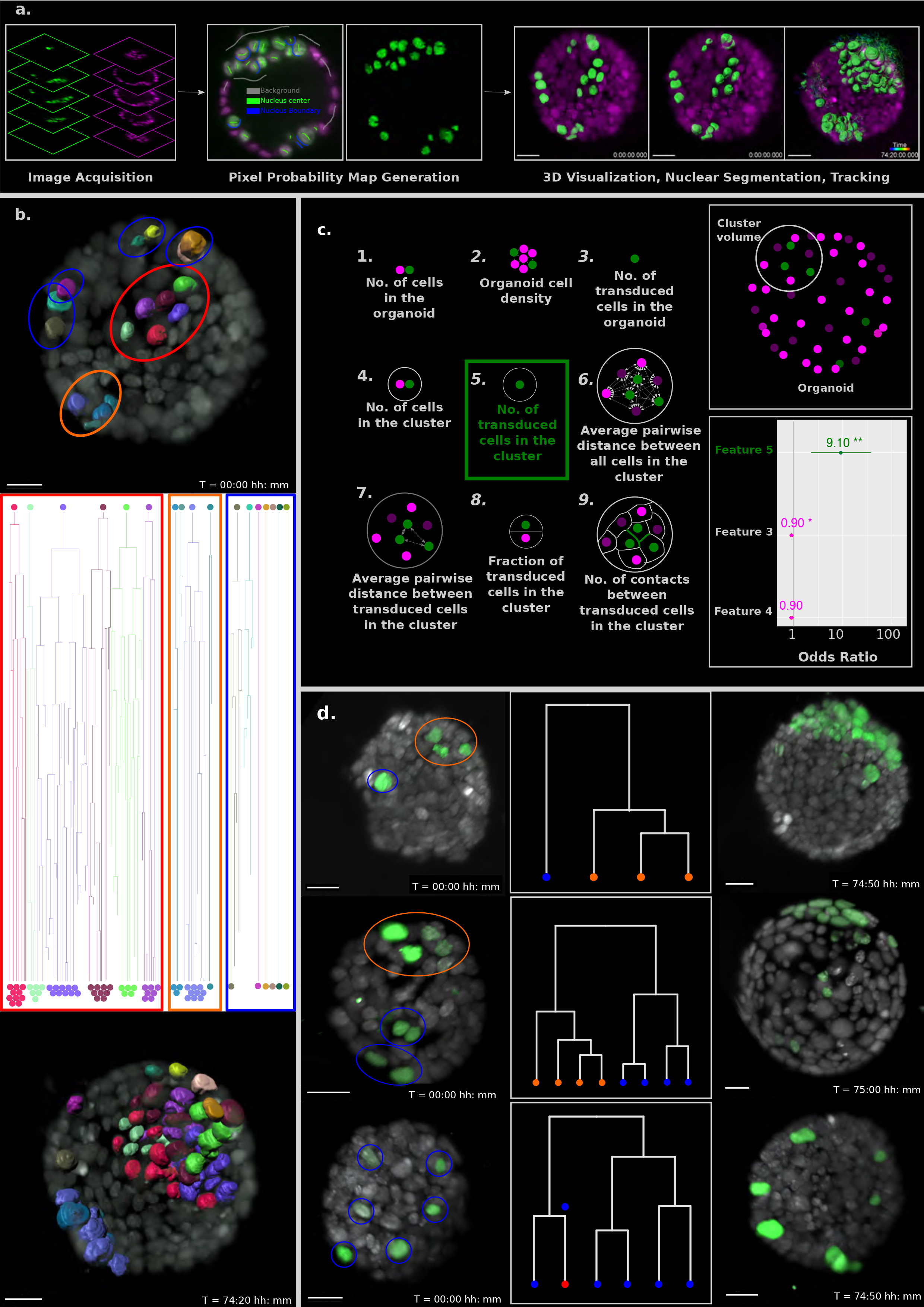
Proximity of transformed cells in a normal epithelium enhances tumor proliferation and establishment. **(a)** Schematic representation of the big-image data analysis pipeline developed to analyze the light sheet microscopy images. Images are acquired in two channels (H2B-mCherry in magenta and H2B-GFP in green) at 10-minute intervals for 3-4 days. Big Data Processor Fiji plugin is used to pre-process the raw images and CATS Fiji plugin is used for generation of pixel probability maps (Supplementary Fig.4). Image pixels of the H2B-GFP images are classified into background (black), nucleus centre (green), nucleus boundary (blue) classes by manual training. Processed raw images along with the probability maps from the nucleus center channel (green) are exported to Imaris for 3D visualization, nuclear segmentation and single cell tracking. **(b)** Single cell tracking results for every cell in a representative B organoid transduced with the Inducer-reporter virus. Top panel shows the organoid at the beginning of the time-lapse (24 hours post induction) with each transduced cell surface rendered with Imaris. The middle panel shows the lineage trees of each individual cell over the time lapse recording. Lineage trees of single cells are grouped into proliferative (highlighted in red, orange) and non-proliferative (highlighted in blue) cell clusters. The bottom panel shows the organoid at the end of the time-lapse (~76 hours post induction with doxycycline). Color coding of each cell maintained in all panels. Scale bar, 15μm. **(c)** Schematic representation of the 9 features of stochastically transformed cells extracted at the beginning of time lapse imaging. These features were assessed for their impact on tumor cell proliferation within B organoids transduced with the Inducer-reporter virus using logistic regression. Lower right panel: Coefficients (represented as odds ratios) of the three features included in the best logistic regression model, colored horizontal bars represent the 95% confidence interval of the estimate. ** indicates p-value (of having no effect) < 0.01, * indicates p-value <0.05. The vertical grey line indicates the position of no effect. **(d)** Representative B mammary organoids stochastically transduced with the Inducer-reporter virus and induced with doxycycline. Left panels show organoids 24 hours post induction. Color highlights indicate clusters of transduced cells identified from hierarchical clustering (shown in middle panels) with proliferative clusters highlighted in orange and non-proliferative clusters highlighted in blue. Right panels show the same organoids ~72-76 hours post induction. Scale bar, 20μm.

To better understand the parameters that positively affect a transduced cell in the stochastic tumorigenesis model to start proliferating and establishing a tumor within a normal epithelium, we extracted 9 features of the organoids (n=20) and all the transduced cells in these organoids (n=150) at the start of the imaging (Fig.2c, left panel). Following observations that tumors originate from groups of independently transduced oncogene expressing cells, we defined clusters of cells in the stochastic model that contain both oncogene-expressing cells as well as normal cells of the organoid and thereby can be used to ascertain the effect of the immediate microenvironment on tumor cell proliferation. Since all the oncogene expressing cells could be tracked over time, each cluster was associated with either a tumor outcome, or a failure to do so. To identify which features were linked to this outcome, we fitted a logistic regression model that shows that only one feature, the “number of transduced cells in the cluster”, positively drives tumor formation within an organoid. Each additional transduced cell in a cluster increases the odds of this cluster forming a tumor by 9 (Fig.2c, lower right panel). This is further demonstrated by representative organoids shown in Fig.2d where the cells that are likely to proliferate, cluster together at the start of the time lapse imaging and the non-proliferative cells are more sparsely located within the organoid, as verified by the hierarchical cluster analysis.

Our results indicate that a proximity-controlled interaction or signaling network between different transformed cells might be imperative to tumor outgrowth in a normal epithelium. This might be due to the repressive effect that an intact polarized tissue layer exerts on single early stage cancerous cells^18^ or rooted in paracrine effects (e.g.via microRNAs^19^). Indeed, studies on loss of important polarity proteins have highlighted their function as non-canonical tumor suppressors in breast tumorigenesis^20, 21^, however, other reports^22, 23^ cannot confirm these observations which were all obtained from tissue wide transformed model systems. Clearly, to better interrogate deficiencies in cell-cell interactions and to settle such conflicting reports, there is a need for a more detailed analysis, employing a model system that does not show modification of all cells in the tissue.

The interaction of tumor cells with the immediate microenvironment has been subject of extensive studies with regards to immune cells^24^ and other tumor associated celltypes^4^, however, the interaction with the normal neighboring cells has not been explored in real time using an organotypic model system. Here, we have developed an integrated approach that allows us to follow cell fates in the first stochastic breast tumor model of primary cells. The amenability of this system to interference with small molecule inhibitors, viral shRNA vectors and genomic editing has the potential to further our understanding of the mechanisms important during tumor initiation. The ability to distinguish marked tumor cells from the normal epithelium will now allow us to perform single cell RNA sequencing analysis on select sorted cells. This will help delineate the signaling networks within the immediate tumor microenvironment.

Taken together, we strongly believe that our integration of a true stochastic tumor model with the ability to image single-cell fates will successfully bridge the gap between genetically modified model systems and the clinical situation, helping gain novel insights on breast cancer.

## Contributions

M.J. and L.C. conceived the model system idea. L.C. performed the cloning and culture experiments. A.A. performed the imaging and implemented the image analysis workflows. T.C. developed the Fiji plugins and designed the image analysis workflow. S.R., M.W., and M.L. provided support for imaging. J.K.H. performed the computational feature analysis. A.A. and M.J. wrote the manuscript and all authors provided feedback. M.J. supervised the work.

## Acknowledgements

The authors want to thank Sylwia Gawrzak, Ksenija Radic, Rocio Sotillo, Robert Prevedel and Jan Ellenberg for critically reading the manuscript and Marta Garcia Montero for mouse husbandry. This study was technically supported by EMBL Advanced Light Microscopy Facility (ALMF) and the EMBL Laboratory for Animal Resources (LAR).

## Online Material and Methods

### Animals

The mouse strains TetO-MYC/ MMTV-rtTA^1^ and TetO-Neu/ MMTV-rtTA^2^, that have been previously described, were bred in order to establish the tri-transgenic strain TetO-MYC/TetO-Neu/ MMTV-rtTA (T) or bi-transgenic strain TetO-MYC/TetO-Neu (B). Reporter H2B-mCherry was crossed into the B and T lines using a R26-H2B-mCherry line {Abe, 2011 #1294}^3^(RIKEN, CDB0239K). All ten mammary glands were harvested (from virgin female mice between 8-10 weeks old), digested and singularized for establishing organoid cultures. All mice used in this study were housed according to the guidelines of the Federation of European Laboratory Animal Science Associations (FELASA).

Rational for the use of these oncogenes: Her2 is overexpressed in ~20% of breast cancers^4^, MYC in 15-50% of human breast cancer^5^. The combination of Myc and Her2 is found in highly aggressive human breast cancer^6^:and - in fact- Her2 and MYC strongly accelerate tumour onset In the combined transgenic animals (average 45 days) as compared to single transgenic animals (MYC 155 days, Her2 99 days), In all cases tumors regress rapidly to non-palpable state following oncogene silencing.

### Lentivirus cloning and production

The lentivirus design is based on pWPXL backbone, which was a gift from Didier Trono (Addgene #12257). The coding region from the original plasmid was excised using ClaI and NdeI in order to insert a new multiple cloning site (MCS). The pGK promoter was PCR amplified from pLVPT-GDNF-rtTR-KRAB-2SM2, which was a gift from Patrick Aebischer & Didier Trono (Addgene #11647) and cloned using XhoI and EcoRI restriction sites. For the plasmid pLenti-rtTA-GFP the synthetic region rtTA-p2A-H2B-GFP was cloned downstream of the pGK promoter using EcoRI and NheI sites. The plasmid pLenti-Null-GFP is derived from the pLenti-rtTA-GFP by removing the rtTA sequence, using the restriction sites EcoRI and BamHI, and retaining H2B-GFP in the coding region. For production of lentivirus particles, we seeded 1.6 x 10^7^ HEK-293T cells (Lenti-X - Clontech Cat. # 632180) in 500cm^2^ square dishes (Corning Cat. # 431110). After 24 hours, the cells were supplemented with media containing 25uM of chloroquine diphosphate (Sigma-Aldrich Cat. # C6628). After a 5-hour incubation, using 360 *μ*g of polyethyleneimine (4 *μ*g for each *μ*g of plasmid), we transfect the cells with a mixture of endotoxin free plasmids: 20 *μ*g pCMV-VSV-G (Addgene #8454); 30 *μ*g psPAX2 (Addgene #12260); 40 *μ*g transfer plasmids pLenti-rtTA-GFP or pLenti-Null-GFP. We harvested the media after 48 hours, 72 hours and 96 hours after transfection. Concentration of the lentivirus from the collected media was performed using an ultracentrifuge (Beckman Sw32 rotor) at 25,000 rpm for 2h at 4°C. The lentivirus pellet was resuspended in 1000 *μ*l of HBBS buffer, aliquoted and stored at −80°C. The lentivirus titer was measured using FACS analyses as described by Kutner and colleagues^7^.

### 3D organoid cultures

Mammary glands harvested from mice (see above), were digested in order to prepare a single cell solution. For this, the tissue was divided in four loosely capped 50 ml falcon, each supplemented with 5 ml serum-free media (DMEM/F12 supplemented with 25mM HEPES and 1% Pen Strep(100 U/ml Penicillin; 100 *μ*g/ml Streptomycin; ThermoFisher Cat. # 15140122)) and 750 U of Collagenase Type 3(Worthington Biochemical Corp Cat. # LS004183), 20 μg of Liberase (Roche Cat. # 5401020001) and incubated overnight at 37°C and 5%CO_2_. The glands were then mechanically disrupted using a 5 ml pipette, and washed in PBS before being pelleted at 1000 rpm for 5 minutes. The cell pellet was resuspended in 5 ml of 0.25% Trypsin-EDTA and incubated for 45 minutes at 37°C and 5%CO_2_. The enzymatic reaction was then neutralized using 40 ml of serum supplemented media (DMEM/F12 with 25mM HEPES, 1% Pen Strep and 10% FBS Tetracycline Free certified (Biowest Cat. # S181T). The cells were pelleted again, resuspended in Mammary Epithelial Cell Basal Medium (PromoCell Cat. # C-21210) and seeded in collagen coated plates (Corning Cat. # 354400) overnight at 37°C and 5%CO_2_. This allows for epithelial cells to adhere to the surface of the plates while the other cell types float on top in the media and can be easily removed by vacuum suction. The epithelial cells were detached from the collagen coated plates by incubating them with 0.25% Trypsin-EDTA for 5-7 minutes at 37°C and 5%CO_2_, following inactivation with serum supplemented media. The single cell solution was pelleted, resuspended in MEBM and counted. We mixed 50,000 cells with 90 *μ*l of Matrigel Matrix basement Membrane growth factor reduced phenol red free (Corning Cat. # 356231), and seeded this mixture into a 12 well plate (Corning Cat. # 3336) and incubated it for 30-40 minutes until the matrigel solidified. The gels were supplemented with 1.5 ml MEBM and allowed to grow at 37°C and 5%CO_2_.

For transduction, after 3 days of growth, the gels were mechanically disrupted and placed in a 15 ml falcon. Two disrupted gels were placed in one 15 ml falcon with 2ml of MEBM supplemented with 25U of Collagenase type I and 5 *μ*g of Liberase. Following incubation in this solution for 2 hours at 37°C and 5%CO_2_, when the matrigel was totally digested, the organoids were washed 3 times with 15 ml of serum supplemented media and once with 15 ml of serum free media, and pelleted at 1000 rpm for 5 minutes. We then supplemented the organoid pellet (from two original gels) in 10 *μ*l of MEBM and added 6 x 10^5^ lentivirus particles to the solution. We then mixed this solution with 90 *μ*l matrigel and plated it in 35 mm dishes (Greiner Bio-One Cat. # 627160) and placed in incubator for 30-40 minutes until the matrigel solidified. The gels were supplemented with 3 ml MEBM and incubated for 2 days at 37°C and 5%CO_2_ in order to allow for organoid recovery and lentiviral gene expression.

For induction of oncogenes in the cells of the organoids, doxycycline (Sigma Cat. # D9891) was supplemented in the media. 800 ng/ml of doxycycline was used to induce T organoids and 600 ng/ml was used for B organoids. qPCR analysis was used to standardize the doxycycline dosage for B organoids (see below).

### qPCR analysis

The qPCR technique was performed following the MIQE guidelines, where the total RNA was isolated from the mammary gland organoids using RNA PureLink Mini Kit (ThermoFisher Cat. # 12183018A) and 2.5ug was reverse transcribed to cDNA using SuperScript VILO cDNA Synthesis Kit (ThermoFisher Cat. # 11754050). Using Primer3 software we designed specific primers for DNA intercalating fluorescent dye approach for the transgenes Neu (Forward: CGTTTTGTGGTCATCCAGAACG and Reverse: CTTCAGCGTCTACCAGGTCACC) and c-MYC (Forward: GCGACTCTGAGGAGGAACAAGA and Reverse: CCAGCAGAAGGTGATCCAGACT). As endogenous controls, mCherry (Forward: GAGGCTGAAGCTGAAGGAC and Reverse: GATGGTGTAGTCCTCGTTGTG) and Pum1 (Forward: AATGTGTGGCCGGATCTTGT and Reverse: CCCACAGTGCCTTATACACCA) were used. Primer efficiency was verified and established between 95% and 105% Each sample was analyzed in duplicate and non-template controls were used in each qPCR run. Analyses were carried out using a StepOne device (Applied Biosystems, USA). Analysis of relative gene expression data was performed according to the 2-ΔΔCq method and the results were expressed as fold change of ΔΔCq values obtained from the reference T800 organoids (Supplementary Figure 1).

### Immunofluorescence staining

Matrigel cultures were grown as described above and plated on Nunc™ Lab-Tek™ II (Thermo Cat. # 155382) chambers. At pre-defined timepoints, the gels were fixed using 4% PFA for 2-3 minutes, following 3 washes with PBS. The gels were blocked with 10% goat serum for 2 hours at room temperature, followed by incubation with primary antibodies was done overnight at 4°C. The remaining immunofluorescence staining was performed as per standard protocol for c-MYC (Cell Signaling Technologies, Cat. # D84C12, 1:900), alpha6-integrin (Millipore Cat. # MAB1378, dilution 1:80) and ZO1 (Life Technologies Cat. # 61-7300, dilution 1:500). The nuclei were counter stained with 1:1000 DAPI (ThermoFisher Cat. # 62248, 1mg/ml, dilution 1:1000) and mounted in anti-fading mounting medium (VECTASHIELD® Mounting Medium with DAPI (Vecto Cat. # H1500-10)). Please note that the c-MYC antibody (Cell Signaling Technologies, Cat. # D84C12) recognizes specifically the human protein, which is transgenically expressed and does not recognize endogenous mouse MYC protein.

Stained gels were imaged on Leica SP5 confocal microscope using 63x water lens and the LAS AF imaging software.

### Light sheet microscopy

#### Sample holder preparation and mounting

Imaging was performed on the InVi SPIM inverted light-sheet microscope (Luxendo Light-Sheet, Bruker Corporation). Sample mounting for the InVi SPIM is suitable for 3D matrigel cultures that are used to grow and transduce mammary organoids (see above). The sample holder is made of medical grade plastic (PEEK). A 25 μm thin membrane (FEP; Luxendo) with a refractive index matching that of water is glued to the upper surface of a groove in the sample holder with a biocompatible silicone glue (Silpuran 4200; Wacker), forming a trough with transparent bottom (Supplementary Fig. 2). Matrigel cultures were carefully cut with a scalpel into rectangular slivers and transferred onto the FEP membrane’s trough. Once the gel sliver was aligned in place, 20-30 *μ*l of fresh matrigel drops were poured onto the gel sliver in the sample holder until there was a thin layer of liquid matrigel on top of the gel sliver. The setup was incubated for 20 minutes at 37°C in a 5% CO_2_ incubator to allow the matrigel layer on top to solidify. Once the gel was solidified, 600-800 *μ*l of MEBM supplemented with/without doxycycline was added to the sample holder’s FEP sheet trough. Preferably, freshly mounted sample gels were allowed to settle overnight in the incubator to prevent any gel drift during imaging, when the holder is placed into the imaging chamber of the microscope. The imaging chamber acts as an incubator with environmental control and it has a reservoir for immersion medium, which is filled with water so that both objective lenses and the bottom of the sample holder are below the water surface (Supplementary Figure 2).

#### Imaging configuration and conditions

The InVi SPIM is equipped with a Nikon CFI 10x/0.3NA water immersion lens for illumination and a Nikon CFI-75 25x/1.1NA water immersion lens for detection. For excitation of GFP and mCherry, 488 nm and 594 nm laser lines were used, respectively, while emission was selected using a 497-554 nm band pass filter and a 610 nm long pass filter, respectively. 3D image stacks were acquired with a light-sheet thickness of 4 *μ*m, a final magnification of 62.5x, resulting in 104 nm pixel size. The In-Vi SPIM environmental control was set to 37 °C, 5% CO_2_ and 95% humidity. A series of optimization experiments, involving different laser powers, exposure times and z-step sizes yielded laser powers of 13 *μ*W for 488 nm and 36 *μ*W for 594 nm, 100 millisecond exposure time per frame and 1*μ*m z-spacing between frames to be optimal for long term imaging (96-120 hours) without photo-bleaching or photo-toxic effects on growth.

Images were recorded as 2D planes ranging from 100-500 in number, depending on the organoid size. Each 3D stack of planes was recorded in 2 channels - mCherry (all cells) and GFP (transduced cells). Depending on the duration of the time lapse imaging, 450-600 image stacks (equivalent to ~72-96 hours) were recorded per organoid at 10-minute intervals.

### Image Analysis

Big Data Processor^8^, a Fiji plugin for lazy loading of big image data, was used to visualize the images in 2D slicing mode, crop stacks in x, y, z, and t, bin images (3 × 3 × 1 in x, y, z), perform chromatic shift correction between channels and convert .h5 files from the InVi SPIM into an Imaris compatible multi-resolution file format (.ims) for further analysis (Supplementary Fig. 4).

The oncogenic cells (H2B-GFP channel) displayed heterogeneous morphologies as well as varying intensity textures, making it difficult to segment them using conventional thresholding approaches. We thus used a trainable segmentation approach to convert the raw intensity values into pixel probability maps, using the Fiji plugin CATS^9^ (Context Aware Trainable Segmentation). Using the H2B-GFP channel images as input, we trained three pixel classes: background, nucleus center and nucleus boundary. For training we drew about 20(background), 120(nucleus center), 100(nucleus boundary) labels distributed across the different time-frames of the movie. After feature computation and training of a Random Forest classifier the whole dataset was processed on EMBL’s high performance computer cluster. The segmentation of one data set -typically 100 timepoints- is distributed across few hundred jobs, each job using 32 GB RAM, 16 cores, and running for about 30 minutes. The nucleus center probability maps were then exported from CATS and added as an additional channel to the converted intensity data (Supplementary Fig. 4).

The data were then loaded into Imaris^10^ for 3D visualization and further processing. Using the Imaris’ Surfaces function, we segmented the nucleus center probability maps into objects. To do so, probability maps were manually thresholded, using a surface smoothening parameter of 0.3 *μ*m; the minimum quality parameter for seed points was set to 0.1, and object splitting was applied for objects larger than 5.5 *μ*m. Objects with volumes less than 20 *μ*m^3^ were excluded. Next, all objects were tracked over time using Imaris’ Lineage tracking algorithm with a maximum distance between objects in subsequent time-points limited to 10 *μ*m and a maximum gap size between identification of the object in a particular track limited to 10 time points. Analysis of segmentation results is shown in Supplementary Figure 5. Most of the errors in the object segmentation were false merges, where two cells were segmented as one. This kind of error is frequently not sustained in the previous or following time-points and the maximum gap size parameter of the tracking algorithm thus frequently provides correct tracks nonetheless. The resulting lineage trees of proliferating tumour cells within the organoid were corrected manually within Imaris, e.g., excluding apoptotic cells and auto-fluorescent debris. Center of mass coordinates of each cell were measured and exported from Imaris for subsequent feature analysis (Figure 2c).

### Feature Analysis

Observations suggest that tumors in organoids originate from clusters of oncogene-expressing cells produced by independent transduction events. To identify these clusters, we computed the pairwise Euclidean distances between all oncogene-expressing cells in an organoid at the start of the experiment and applied hierarchical clustering with complete linkage. Clusters were identified automatically by cutting the branches of the trees using the dynamic tree cut algorithm^11^. This defined a cluster as a group of oncogene-expressing cells that are closer to each other than to other oncogene-expressing cells of the same organoid. Note that a cluster can be composed of a single cell if this cell is comparatively isolated from other transduced cells. For each cluster we identified the following features as possibly linked to tumor formation: (1) number of cells in the organoid (2) cell density expressed as the ratio of number of cells to organoid surface area computed by assuming the organoid is a sphere with diameter equal to the distance between the two most distant cells (3) number of oncogene-expressing cells in the organoid (4) number of cells (including both oncogene-expressing and normal cells) in the cluster volume defined as the sphere centered at the center of mass of the cluster with diameter equal to the distance between the two farthest oncogene-expressing cells of the cluster (5) number of oncogene-expressing cells in the cluster (6) average pairwise distance between all cells in the cluster volume (7) average pairwise distance between oncogene-expressing cells in the cluster (8) fraction of oncogene-expressing cells in the cluster volume (9) number of contacts between oncogene-expressing cells in the cluster. Two cells are presumed in contact if they are less than the average cell diameter + 2 standard deviation apart.

Oncogene-expressing cells were tracked over time and a cluster was associated with a tumor outcome if any of its cells lead to tumor formation. To identify which features were linked to this outcome, we took an information-theoretic approach to model selection. We fitted a logistic regression model for all possible linear combinations of features and selected the best model based on the Akaike information criterion (with correction for small sample sizes)^12^ (Ref 9). This model included only three features: number of oncogene-expressing cells in the cluster, number of oncogene-expressing cells in the organoid and number of cells in the cluster of which only the first (number of oncogene-expressing cells in the cluster) contributed significantly to tumor formation with an odds ratio of 9.1 (Figure 2c). Computing relative variable importance across all models also indicated that the number of oncogene-expressing cells in a cluster is the most important feature.

