## Supplementary Figures for "Tracking the cells of tumor origin in breast organoids by light sheet microscopy"

### Supplementary Figure 1

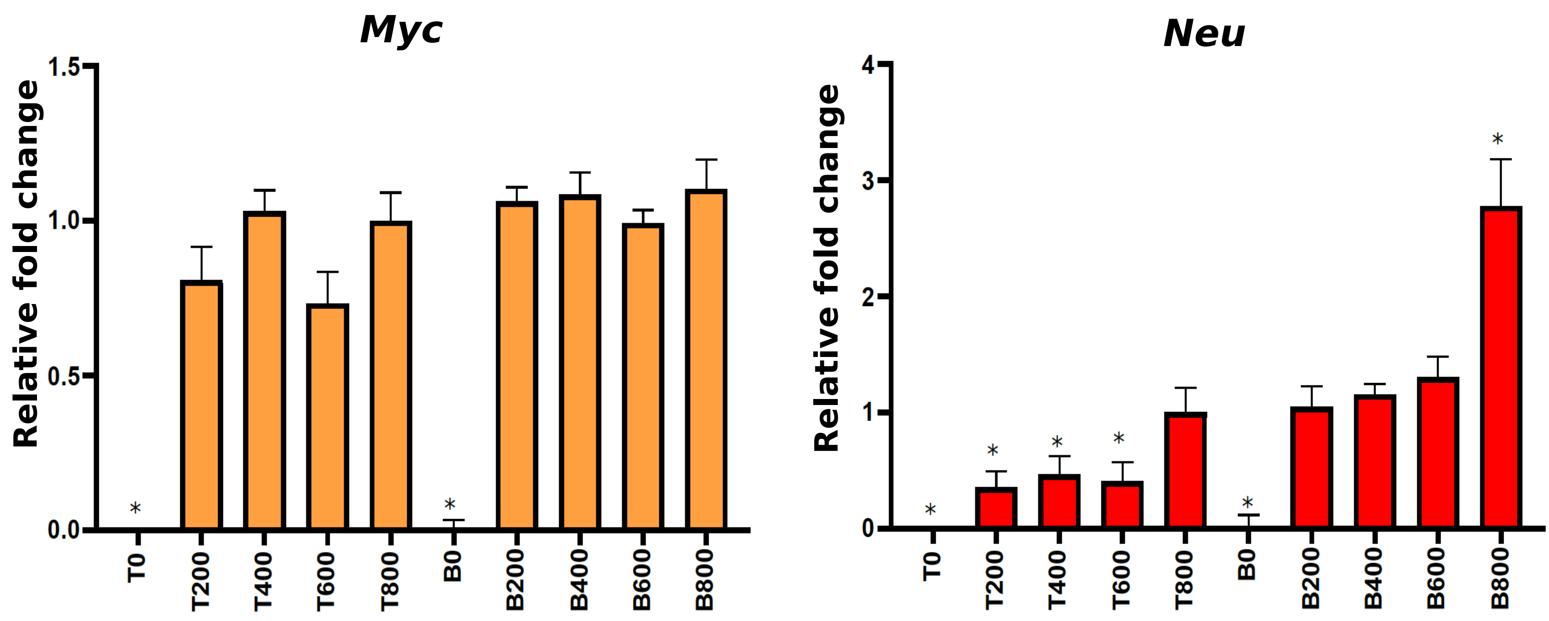


**Supplementary Figure 1. --**
**Normalization of doxycycline concentration for the stochastic model using qPCR analysis**
Fold changes in the mRNA expression of transgenes, *Myc* and *Neu* in transduced mammary epithelial cells of B mice (n=2) infected with Inducer-reporter virus or T mice (n=2) with Reporter virus. The doxycycline dosage of 800 ng/ml (T800) is well established in the T cells and was used as control to normalize the gene expression, and also to determine the dose for transduced B cells (600 ng/ml). Data represented as mean ± SEM; *P <0.05.

### Supplementary Figure 2

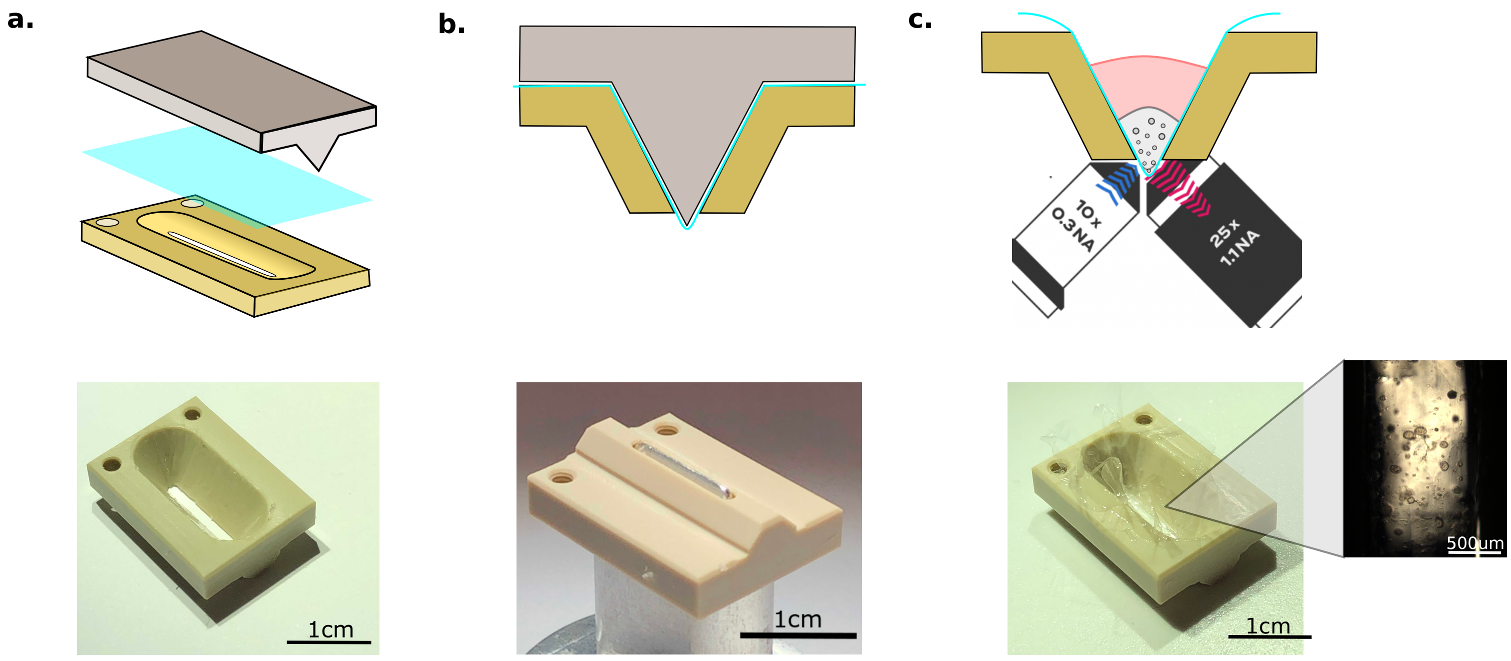


**Supplementary Figure 2 --**
**Sample holder preparation and sample mounting**
The FEP membrane is glued onto the sample holder (a) with the help of a mold and biocompatible glue(b). Gel slivers are transferred to the FEP sheet trough in the sample holder (c) and overlaid with fresh matrigel to prevent drift during imaging. Media is added after the upper matrigel layer solidifies.

### Supplementary Figure 3

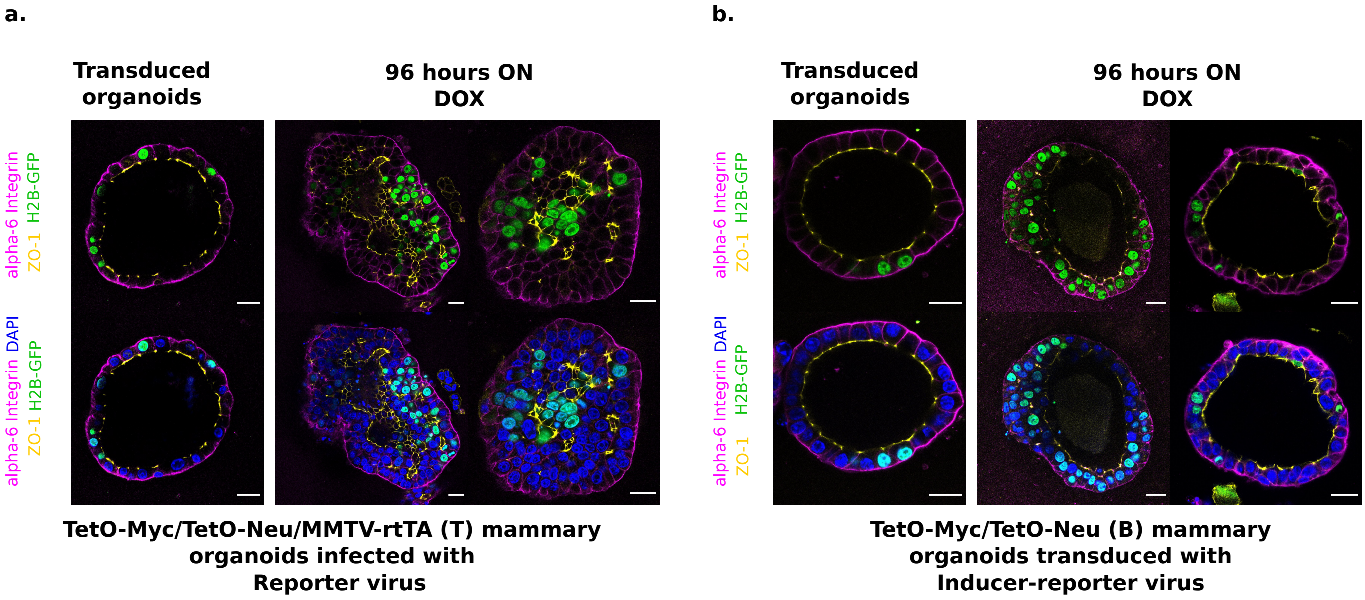


**Supplementary Figure 3--**
**Immunofluorescence staining of oncogene induced organoids shows dual phenotype in stochastic model**
Representative immunofluorescence staining images of PFA (4%) fixed 3D gels with **(a)** T organoids (transduced with Reporter virus) and **(b)** B organoids (transduced with Inducer-reporter virus), before induction and 96 hours post induction with doxycycline. Polarity markers include alpha-6-Integrin (magenta) and ZO-1 (yellow). Transduced cells are marked with GFP (green) and nucleus is counterstained with DAPI (blue). Scale bar, 20 µm.

### Supplementary Figure 4

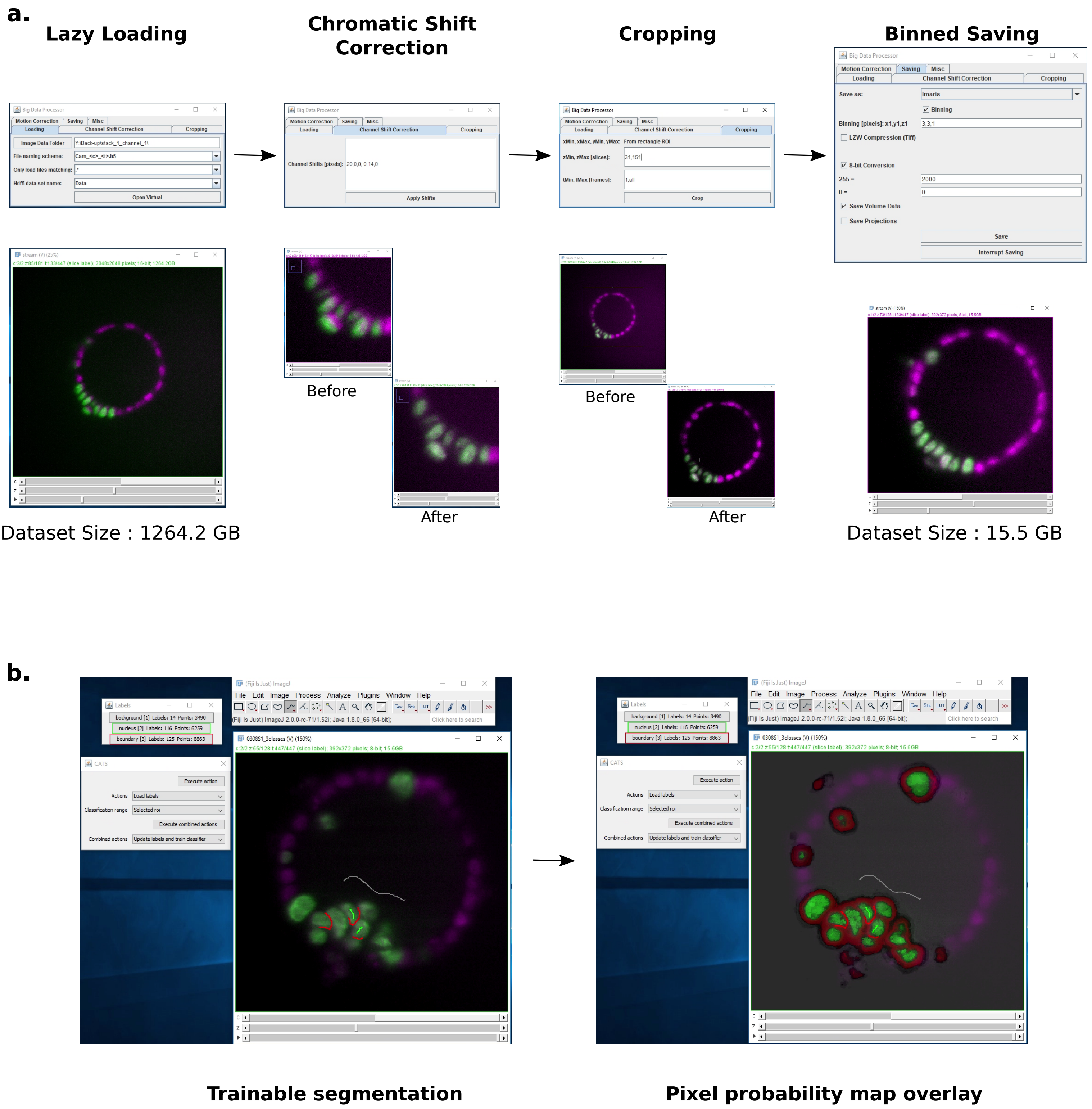


**Supplementary Figure 4--**
**Image pre-processing and pixel probability generation using Fiji plugins**
**(a)** The Big Data Processor Fiji plugin was employed for pre-processing light-sheet microscopy images. Two-channel raw images were lazily loaded in 2D slice mode for visual inspection. Channel shift correction was performed to align the two channels. Then, the whole dataset was cropped in x,y,z to remove black pixels and empty planes. The cropped dataset was then saved in an 8-bit Imaris format with 3x3 binning applied in x and y. **(b)** The CATS Fiji plug-in was used to generate pixel probability maps for H2B-GFP images. Left panel shows the manual training done by drawing labels on the dataset to classify pixels into 3 classes - background (grey), nucleus boundary (red) and nucleus center (green). The right panel shows the pixel probability output for all three classes overlaid on the intensity data. Only the pixel probabilities from the nucleus center class were exported from CATS and linked to the Imaris data set for further segmentation and tracking on Imaris.

### Supplementary Figure 5

H2B-mCherry
True Positives
False Merges
False Splits

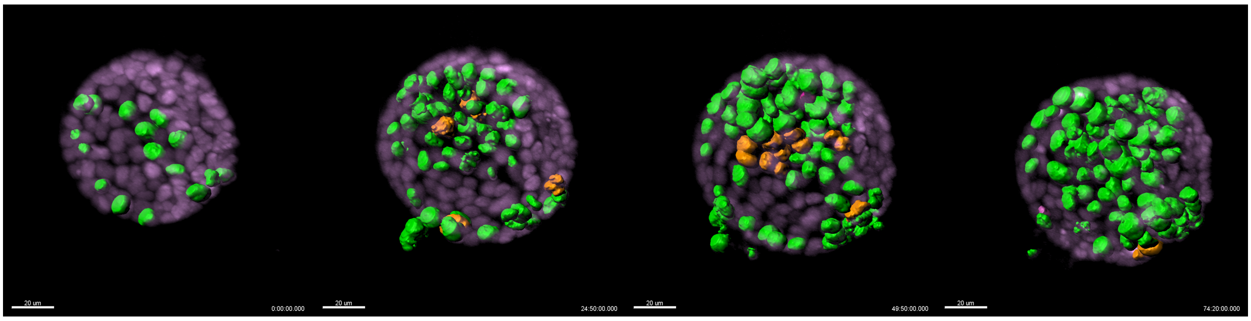

| | T = 1 | T = 150 | T = 300 | T = 447 |
|---|---|---|---|---|
| **True Positives** | 17 | 52 | 69 | 74 |
| **False Merges** | 0 | 4 | 6 | 1 |
| **False Splits** | 0 | 0 | 2 | 0 |
| **False negatives** | 0 | 1 | 0 | 0 |
| **Ground Truth** | 17 | 61 | 82 | 76 |

**Supplementary Figure 5--**
**Cell segmentation accuracy**
Image panels show the H2B-mCherry signal (magenta) along with the segmented H2B-GFP cells of an organoid at four equidistant timepoints. Segmentation accuracy was assessed, counting: True Positives (correctly segmented cells, highlighted in green), False Merges (two cells merged as one, highlighted in orange), and False Splits (one cell split in two, highlighted in purple). Unidentified cells are indicated as False Negatives and Ground Truth indicates the actual number of cells at each timepoint. The average True Positive to Ground Truth ratio for the four time-points is 0.92. Scale bar, 20 µm.
